# Ag nanoparticles-based antimicrobial polycotton fabrics to prevent the transmission and spread of SARS-CoV-2

**DOI:** 10.1101/2020.06.26.152520

**Authors:** Guilherme C. Tremiliosi, Luiz Gustavo P. Simoes, Daniel T. Minozzi, Renato I. Santos, Daiane C. B. Vilela, Edison Luiz Durigon, Rafael Rahal Guaragna Machado, Douglas Sales Medina, Lara Kelly Ribeiro, Ieda Lucia Viana Rosa, Marcelo Assis, Juan Andrés, Elson Longo, Lucio H. Freitas-Junior

## Abstract

Pathogens (bacteria, fungus and virus) are becoming a potential threat to the health of human beings and environment worldwide. They widely exist in the environment, with characteristics of variety, spreading quickly and easily causing adverse reactions. In this work, an Ag-based material is used to be incorporated and functionalized in polycotton fabrics using pad-dry-cure method. This composite proved to be effective for inhibiting the SARS-CoV-2 virus, decreasing the number of replicates in 99.99% after an incubation period of 2 minutes. In addition, it caused 99.99% inhibition of the pathogens *S. aureus*, *E. coli* and *C. albicans*, preventing cross-infections and does not cause allergies or photoirritation processes, demonstrating the safety of its use.

## INTRODUCTION

Pathogenic microbes are becoming a potential threat to the health of human beings and environment worldwide. Today, the humanity has experienced epidemic diseases caused by both new and well known viruses, including hepatitis C, HIV/AIDS, SARS-CoV, MERS, Lassa fever, Zika virus, and Ebola virus, as well as Yellow Fever, Influenza, and Measles virus, which are more widespread but can be severe, despite the availability of vaccines.[1] Severe acute respiratory syndrome coronavirus 2 (SARS-CoV-2) is a novel coronavirus that causes the coronavirus disease 2019 (COVID-19). Since its first detection in December 2019,[2] it has affected millions of people worldwide, carrying a mortality rate much higher than the common flu. These public health outbreaks driven by emerging COVID-19 infectious diseases constitute the forefront of global safety concerns and significant burden on global economies. While there is an urgent need for its effective treatment based on antivirals and vaccines, it is imperative to explore any other effective intervention strategies that may reduce the mortality and morbidity rates of this disease.

In the absence of an effective vaccine, it is expected that not only the current pandemic will continue for several months, but other outbreaks caused by SARS-CoV-2 may take place in the future, in the coming months or years.[3] Furthermore, unknown viruses and/or pathogens will likely emerge again, and their pathogenicity, spread, contagion, and mechanism of action will be inquired. This current global crisis alarms us to the fact that we urgently need to prepare ourselves for a new and unpredictable epidemic in the future.

The human body is a diverse ecosystem that harbors hundreds of trillions of microbes (bacteria, fungi and viruses)[4] that might be or become pathogenic under certain circumstances. The development of innovative materials capable of avoiding the transmission, spread, and entry of these pathogens into the human body is currently in the spotlight. Highly effective agents are needed not only to control the emergence of new COVID-19 pandemia, their increased proliferation capability, and resistance that severely impact public health, but it is also fundamentally essential to explore strategies for the preparation and application of new materials against pathogen infection.

Often, the surface of a material is the medium by which the human body interacts with microbes. Therefore, anti-pathogen strategies based on chemical modification of the material surface have been developed. One procedure is to form a layer on the surface of the material, thereby reducing the chance of contact between the pathogen and the surface of the material. This greatly reduces the number of pathogens adhering to the surface. Another strategy is that killing the adhered pathogen directly by decorating the surface of the material with the biocide agent.

Use of personal protective equipment is considered to be one of the most important strategies for protecting from transmissible pathogens, particularly when aerosol transmission occurs and when no effective treatment or prophylaxis is available for the disease provoked by these pathogens in question. In the case of COVID-19, for instance, the WHO has recently issued recommendation for widespread use of face masks as an important tool in the control of SARS-CoV-2 spread.[5] Therefore, the current worldwide public health crisis of COVID-19 has highlighted the particularly emergent need for materials that inactivate enveloped viruses on contact for preventing transmission.

Inorganic biocide surfaces and materials have attracted much attention due to their better stability and safety as compared with organic reagents for preventing infections and transmission. Among inorganic agents, silver cation and metal are most widely used. However, Ag cations tend to react with Cl^−^, HS^−^, and SO_4_^2−^ in aqueous solution, forming precipitates, thus losing their biocide activity, which affects the practical application of Ag-loaded biocide agents to a certain extent. Ag and its compounds have been widely used since from ancient time, in 1000 BC, to prevent bacterial growth and wound infections, and make water potable.[6] Ag metal is a precious metal and is easily discolored under light and heat, but for many years, it was used as medical treatment as a broad-spectrum antibacterial compound before the discovery of antibiotics in the early 20^th^ century.[7,8]

Nanotechnology is capable of modifying both Ag cation and metal into their nano range, which dramatically changes their chemical and physical properties. Ag nanoparticles (AgNPs) acquire special attention due to its specificity and environment friendly approach with a wide application in industry and medicine due to its antibacterial, antifungal, larvicidal and anti-parasitic characters. The use of AgNPs has been greatly enhanced due to the development of antibiotic resistance against several pathogenic bacteria, and they are employed in biomedical industry as coatings in dressings, in medicinal devices, in the form of nanogels in cosmetics and lotions, etc.[6,9,10]

According to the literature, there are plenty of protocols focused on the production of hybrid/composite materials based on Ag NPs, whose architecture is driven by different synthetic methods and reaction mechanisms.[11–17] While the precise reasons for this unique chemistry and physics are unknown, the observed structures, their reproducibility, and synthetic control this reaction offers, there is plenty of room to find innovative possibilities for new technologies.[18,19]

Ag NPs have been proven to be most useful because they have excellent antimicrobial properties against lethal viruses, microbes/germs, and other microorganisms. These NPs are certainly the most extensively utilized material among all. Thus, it has been used as antimicrobial agent in different textile industries.[20] The noble metal NPs are considered as more specific and multipurpose agents with a diversity of biomedical applications considering their use in extremely sensitive investigative assays, radiotherapy enhancement, gene delivery, thermal ablation, and drug delivery. These metallic NPs are also considered to be nontoxic in case of gene and drug delivery applications. Thus, metallic NPs can offer diagnostic and therapeutic possibilities simultaneously.[16].

The purpose of this work is to present an innovative material with high bactericide, fungicide and virucide efficiency in their incorporation and application in textile applications such as cotton-based materials that make special biopolymer hosts for composite materials. Finally, it is important to study the reliability of sintered AgNPs, to test and analyze its allergic response, dermatological photoirrritant and photosensitive effects, as well as their antimicrobial, fungicide and antiviral activity

## EXPERIMENTAL: MATERIALS AND METHODS

### Application of chemical finishing onto fabrics

A fine-medium weight 67% polyester / 33% cotton woven fabric (plain weave, 120 g/m²; width 1,60m; ends 35/cm; picks 26/cm; yarn Ne 36 67%Polyester / 33% cotton) purchased from local suppliers (São Carlos/SP, Brazil) was used for the application purpose. An AgNP colloidal solution (AgNP-CS) and an AgNP colloidal solution stabilized with organic polymers (AgNP-OP) were applied on the polycotton fabric using pad-dry-cure method (NanoxClean® Ag+Fresh 5K, and NanoxClean® Ag+Fresh Hybrid, respectively, provided by Nanox Tecnologia S.A. – São Carlos/SP - Brazil). An acrylic-based binder compound was used in the impregnation solution as well (Starcoat Denim 50GL provided by Star Colours LTDA. – Americana/SP, Brazil).

The polycotton fabric cut to the size of 30×30 cm was immersed in the solution containing 5% (% weight basis) of the antimicrobial products and 6% of the acrylic-based binder (% weight basis), for 5 minutes and passed through a laboratory scale padder, with a 72% wet pick-up maintained for all the treatments. After drying (80°C, 3 min) the fabric was annealed at 170°C for 3 min, then washed with deionized water and then dried at 80°C for 3 min in a ventilated oven. All samples were then conditioned at 25°C and 65% relative humidity for 48h. Samples were produced according to Table 1.

**Table 1.** Identification of the polycotton samples.

| Sample Identification | Concentration of AgNP-CS in impregnation bath (% weight) | Concentration of AgNP-OP in impregnation bath (% weight) | Concentration of acrylic-based binder in impregnation bath (% weight) |
| --- | --- | --- | --- |
| Non-Treated Polycotton | - | - | - |
| Polycotton AgNP-CS | 5% | 0 | 6% |
| Polycotton AgNP-OP | 0 | 5% | 6% |

### Characterization

Micro-Raman spectroscopy was performed using an iHR550 spectrometer (Horiba Jobin-Yvon, Japan) coupled to a charge-coupled device (CCD) detector and an argon-ion laser (Melles Griot, United States) operating at λ = 514.5 nm and 200 mW. The spectra were carried out bin the range of 100-3500 cm^−1^. Morphologies of the composites were analyzed by Field Emission Scanning Electron Microscopy (FE-SEM) on a FEI instrument (Model Inspect F50) operating at 1 kV. Fourier Transform Infrared Spectroscopy (FTIR) was performed using a Jasco FT/IR-6200 (Japan) spectrophotometer operated in absorbance mode at room temperature. The spectra were carried out in the range of 400-4000 cm^−1^.

### Assessment of Antimicrobial Activity

The AATCC 147 Parallel Streak Standard Method[21] was used as a qualitative method to evaluate antibacterial activity of the treated fabrics. Sterile plate count agar was dispensed in petri plates. 24 hours broth cultures of the test organisms (*Escherichia Coli* (*E. coli* - ATCC8739) and *Staphylococcus aureu*s (*S. aureus* - ATCC6538) were used as inoculums. Using a 10μL inoculation loop, 1 loop full of culture was loaded and transferred to the surface of the agar plate by making 7.5cm long parallel streaks 1 cm apart in the center of the plate, refilling the loop at every streak. The test specimen was gently pressed transversely, across the five inoculums of streaks to ensure intimate contact with the agar surface. The plates were incubated at 37°C for 18-48 hours. After incubation, a streak of interrupted growth underneath and along the side of the test material indicates antibacterial effectiveness of the fabric.

The quantitative antimicrobial activity assessment of the treated polycotton fabrics was determined according to AATCC Test Method 100[22]. Fabric specimens (circular swatch 4.8 cm in diameter) were impregnated with 1.0 mL of inoculum in a 250 mL container. The inoculum was a nutrient broth culture containing 2.0~3.0 · 10^5^/mL colony forming units of microorganisms. *E. coli* and *S. aureus* were used as a reference for gram-negative and gram-positive bacteria, respectively, and *C. albicans* (ATCC 10231) as a reference for fungus. The microorganisms counted on the treated polycotton fabric and those on a controlled sample were determined after a 24-hour incubation period at 37°C. The antimicrobial activity was expressed in terms of percentage reduction of the microorganism after contact with the test specimen compared to the number of microbial cells surviving after contact with the control. The results are expressed as percent reduction of microorganisms by Eq. (1).

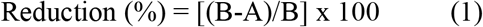

where A and B are the numbers of bacteria or fungus recovered from the antimicrobial-treated and untreated polycotton fabrics in the jar incubated over the desired contact period, respectively.

### Assessment of Antiviral Activity

An adaptation of ISO 18184 Determination of antiviral activity of textile products Standard Method[23] was used as a reference for a quantitative method to evaluate the treated polycotton’s ability to inactivate the SARS-CoV-2 virus particles (SARS-CoV-2/human/BRA/SP02cc/2020 - MT350282), under the tested conditions, at two different time intervals (2 and 5 minutes of contact time). The virus was inoculated into liquid media containing no fabric, treated and non-treated polycotton samples and incubated for 2 different time periods. Then, they were plated onto tissue cultures of Vero CCL-81 cells. After the incubation, the viral genetic material was quantified in each condition using real-time quantitative PCR, and based on the control samples, the ability of each sample to inactivate SARS-CoV-2 was determined.

Briefly, Vero CCL-81 cells were plated onto 24-well plates at 1 × 10^5^ cells per well. The cells were maintained in DMEM high glucose culture medium (Sigma-Aldrich, 51435C) supplemented with 10% fetal bovine serum, 100 U/mL of penicillin, and 100 μg/mL of streptomycin. The plate was incubated at 37 °C, 5% CO_2_ atmosphere for 24 h. Following this period, the medium was removed and replaced with 666.7 μL of DMEM High Glucose/well without supplementation.

Three test specimens, non-treated polycotton control and Ag-based antimicrobial treated polycotton samples, measuring 6.25 cm^2^ apiece, were tested. Each test specimen was placed into a different tube and 1.33 mL of DMEM high glucose medium without supplementation was added to each tube. In parallel, 500 μL of culture medium containing SARS-CoV-2 was diluted in 4.5 mL of DMEM medium without supplementation, and then 333.4 μL of this viral suspension was added to each of the tubes containing the pieces of cloth. The mixtures were incubated with the virus for 2 min and the tubes were homogenized every 30 seconds. After this period, 166.7 μL of each sample was transferred to different wells of the plates containing the cells previously seeded. After a total of 5 min of incubation, an additional 166.7 μL-aliquot was removed from each tube and incubated in other wells on the same plate. As control, the viral suspension was incubated in media without supplementation, with samples collected at 2 and 5 min used to infect Vero cells on the same plate

The plate was incubated for 2 h at 37 °C, 5% CO_2_ for viral adsorption, and after this period, 166.6 μL of DMEM High Glucose medium containing 12% fetal bovine serum were added to each well, making to a final volume of 1 mL of medium/well containing 2% serum. Immediately after adding the medium, the plate was further incubated at 37 °C, 5% CO_2_. After 48 h, the plate was removed from the incubator and 100 μL of the medium from each well (each well a different condition) was removed and placed in lysis buffer to proceed with the viral RNA extraction. For the extraction, the MagMAX™ CORE Nucleic Acid Purification Kit (Thermo Fisher) was used, following the manufacturer’s instructions, on the semi-automated platform MagMAX Express-96 (Applied Biosystems, Weiterstadt, Germany). The detection of viral RNA was carried out using the AgPath-ID One-Step RT-PCR Kit (Applied Biosystems) on an ABI 7500 SDS real-time PCR machine (Applied Biosystems), using a published protocol and sequence of primers and probe for E gene.[24] The number of RNA copies/mL was quantified by real-time RT-qPCR using a specific in vitro-transcribed RNA quantification standard, kindly granted by Christian Drosten, Charité - Universita◻tsmedizin Berlin, Germany, as described previously.[25] The viricidal activity, or viral inactivation, was determined as a percentage related to the control (media without fabric specimen).

The experiment was repeated using the same experimental conditions, but with media incubated with two pieces of test specimens (instead of one) per condition.

### Assessment of Allergic Response

A Human Repeat Insult Patch Test (HRIPT) was performed to determine the absence of the potential for dermal irritability and sensitization of the treated fabrics. The study was carried out in maximized conditions, in which semi-occlusive dressings containing the investigational product and controls were applied to the participants’ backs. The application of the study dressings occurred for six weeks, with three weeks of application alternately, two weeks of rest and a new application of the dressing containing the product in virgin area in the sixth week (challenge). The readings of the application site were performed at each dressing change according to the reading scale recommended by the International Contact Dermatitis Research Group (ICDRG)[26]. Dermatological evaluations were carried out at the beginning and end of the study, and a physician was available for evaluation and assistance to the participants in case of positive or adverse reaction. Participants of both genders, with phototypes III to IV (Fitzpatrick),[27] aged between 21 and 70 were selected. The selected participants were distributed as shown in the Table 2.

**Table 2.** Distribution of selected participants for the HRIPT.

| Evaluation Test | Number of Participants | Gender |  | Age |  |
| --- | --- | --- | --- | --- | --- |
|  |  | Female | Male | Minimum | Maximum |
| Primary Dermal Irritability | 51 | 40 | 11 | 21 | 70 |
| Accumulated Dermal Irritability | 51 | 40 | 11 | 21 | 70 |
| Dermal Sensitization | 51 | 40 | 11 | 21 | 70 |

### Assessment of Dermatological Photoirrritant and Photosensitive Potential

Since exposure to solar radiation can trigger or aggravate adverse reactions to topical products, knowing the behavior of the product on human skin stimulated with ultraviolet radiation is of fundamental importance for proof of safety. Therefore, a unicentric, blind, comparative clinical study to assess the photoirritating and photosensitizing potential was also conducted, with the aim of proving the absence of the irritating potential of the product applied to the skin when exposed to ultraviolet radiation. The study was carried out with dressings containing the product, applied to the participants’ skin and, after removal, controlled irradiation with a spectrum of ultraviolet radiation emission was performed. Readings were performed according to the reading scale recommended by the ICDRG. The study with the participants lasted for five weeks, covering 3 phases: induction, rest and challenge. Dermatological evaluations were performed at the beginning and end of the study, or when there was an indication of positivity or adverse reaction. Participants of both genders, with phototype III (Fitzpatrick), aged between 21 and 62 were selected. The selected participants were distributed as shown in the Table 3.

**Table 3.** Distribution of selected participants for the photoirritating and photosensitizing clinical study.

| Evaluation Test | Number of Participants | Gender |  | Age |  |
| --- | --- | --- | --- | --- | --- |
|  |  | Female | Male | Minimum | Maximum |
| Photosensitization | 25 | 20 | 05 | 19 | 62 |
| Photoirritation | 25 | 20 | 05 | 19 | 62 |

## RESULTS AND DISCUSSION

There are numerous ways to functionalize a textile substrate, from the development of new structures up to finishes that modify the material’s surface. The superficial modification through the incorporation of nanoparticles has been extensively studied and shows potential for obtaining devices with microbicidal activity.[28–33] In this scenario, nanoparticles can advantageously replace micrometric particles used in the finishes to obtain functional fabrics, because it has a greater surface area, resulting in a better adhesion to fabrics and, consequently, greater durability of functionality. Furthermore, it is possible to achieve a pronounced effect with small amounts of material not altering the original properties of the fabric. Traditionally, the pad-dry-cure is the most common finishing route applied to impart different finish treatments on textile fabrics.[34,35] In this way, the interactions between the polycotton fabric and the Ag NPs were investigated, in order to observe how these changes mirrored the microbicidal properties in the new Ag-based fabric.

In order to analyze the local structural order/disorder caused by the addition of Ag-based antimicrobials to polycotton, micro-Raman analyzes were performed. The results are presented in Figure 1. The adhesion and durability of a superficial change in fabrics depends on the surface chemical properties.[28] Since the main component of polycotton is cotton (formed by glucose monomers) and polyester (polyethylene terephthalate), the vibrations refer to C, O and H bonds. The cotton Raman spectra of the samples can be divided into four blocks related to the glycosidic ring skeleton, to OH groups, the CH and CH_2_ groups and acetylation of cotton. The glycosidic ring presents the fingerprint of the cotton Raman spectra. These modes can be observed at 859, 1104, 1123 and 1183 cm^−1^ and represent the symmetrical stretching of C-O-C in the plane, asymmetric and symmetrical stretching of C-O-C in the glycosidic link, and asymmetric stretching of C-C ring breathing.[36] The presence of OH groups in the samples is observed by the modes located at 287 and 1464 cm^−1^, related to the twisting and deformation of the C-OH bonds.[37,38] Four modes related to CH_2_ deformations and twisting are observed in 1000, 1291, 1372 and 1418 cm^−1^. There is still a mode located at 289 cm^−1^ regarding the twisting of the C-CH bond.[39] The acetylation of the cotton used is further confirmed by the modes located at 705 and 795 cm^−1^, referring to deformation O-C=O and the stretching of the H_3_C-C bonds.[40] Like cotton, polyester has its fingerprint given by the modes referring to its aromatic ring and its esters. It is observed in 1613 cm^−1^ the mode referring to the stretching of the C^1^-C^4^ carbon of the aromatic polyester ring, as well as its CH stretching in 3078 cm^−1^.[41,42] It is also possible to observe in 1730 cm^−1^ the stretching of the C=O bonds of the esters and the stretching of the CH bonds of the methyl groups external to the ring, in 2975 cm^−1^.[41–43] It is observed that the addition of Ag-based antimicrobials do not cause significant changes in the polycotton structure at short-range.

**Figure 1.**
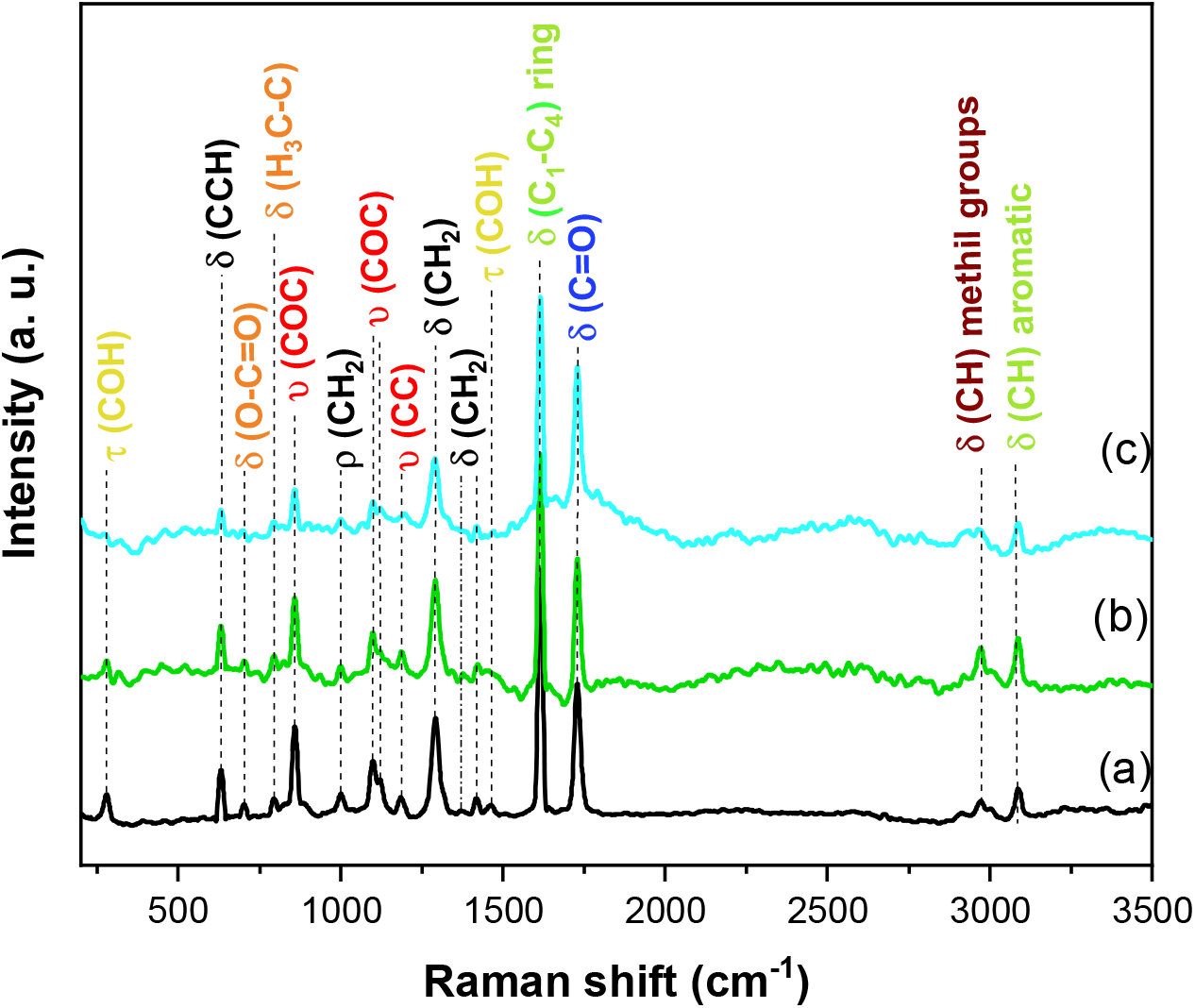
Micro Raman spectra of (a) Non-Treated Polycotton, (b) Polycotton AgNP-CS and (c) Polycotton AgNP-OP.

As a complementary analysis to micro Raman spectroscopy, FTIR measurements were performed to investigate the functional groups of products after the incorporation of the two Ag-based antimicrobial solutions into the polycotton (Figure 2). It is observed in the samples of pure polycotton and those modified with the Ag-based antimicrobials the peaks located at 3540, 2936, 2124, 1947, 1710, 1338, 1147, 832 and 647 cm^−1^. The peaks located at 3540, 2936 and 2124 cm^−1^ refer to OH stretching and CH deformation respectively, the latter being related to the CH_2_ groups of the cellulose structure.[44,45] The peaks located in 1947, 1710, 1338 and 1147 cm^−1^ correspond respectively to H_2_O adsorbed on the polycotton surface, stretching of the CH bond, asymmetric deformation of the C-O-C groups and were attributed to stretching vibrations of intermolecular ester bonding.[45–47] As in the Raman spectra, the cotton fingerprint can be observed in the FTIR due to the presence of the band located at 832 cm^−1^, referring to the asymmetric stretching of the glycosidic ring, especially the C^1^-O-C^4^ bonds.[46] It is observed for the modified polycotton in relation to the non-treated polycotton the displacement of the band located at 1338 cm^−1^, as well as the decrease of the band located around 647 cm^−1^, referent to in-plane bending of O-H mode from the glycosidic units and deformation of the OH, respectively.[48,49] These shifts, as well as the appearance of new bands in the FTIR spectra of the modified polycotton are due to interactions between polycotton and Ag-based antimicrobial additives.[50–52]

**Figure 2.**
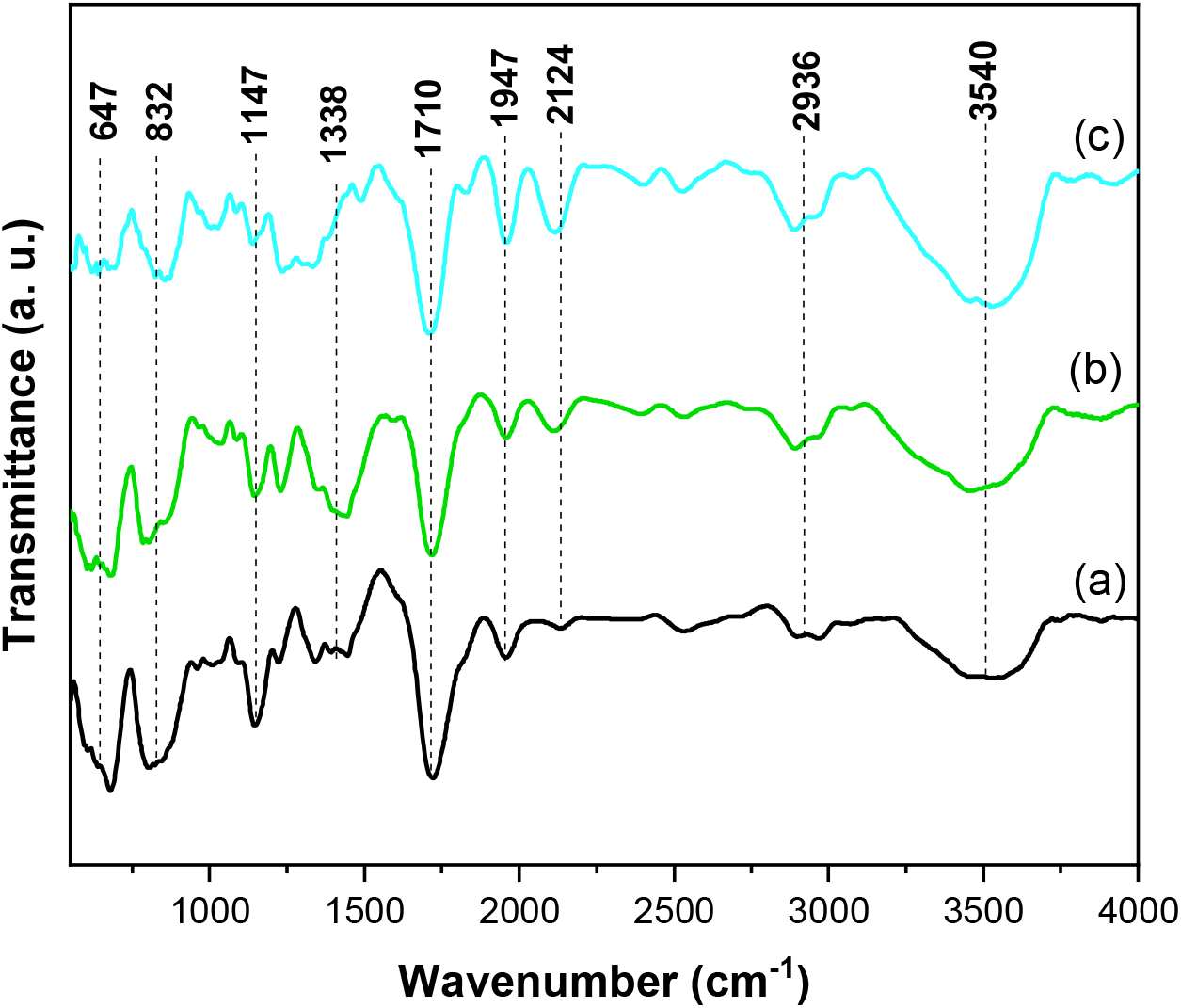
FTIR spectra of (a) Non-Treated Polycotton, (b) Polycotton AgNP-CS and (c) Sample 2.

To observe morphological changes in polycotton fibers, FE-SEM measurements were performed (Figure 3). There are no significant differences in the fiber diameters of the samples, being the average diameter values obtained for the Non-Treated Polycotton, Polycotton AgNP-CS and Polycotton AgNP-OP were 10.62 ± 2.30, 10.22 ± 2.04 and 10.59 ± 2.50 μm respectively. For Polycotton AgNP-CS (Figures 3d-f), it is possible to observe the formation of small Ag nanoparticles on the polycotton surface, with average size of the 23.51 ± 5.18 nm. Similar behavior was obtained by several other authors in works that incorporated AgNPs into polycotton in different ways.[53–60] For Polycotton AgNP-OP, the formation of a smaller amount of Ag nanoparticles with average size higher (126.9 ± 19.5 nm) than the than Polycotton AgNP-CS were observed. In addition, there is a homogeneous distribution of micrometric crystals with well-defined morphology over all polycotton surface fibers, with an average size of 1.62 ± 0.44 μm. These differences are due to the different composition of both Ag-based antimicrobials, which result in different surface effects on polycotton surface fibers.

**Figure 3.**
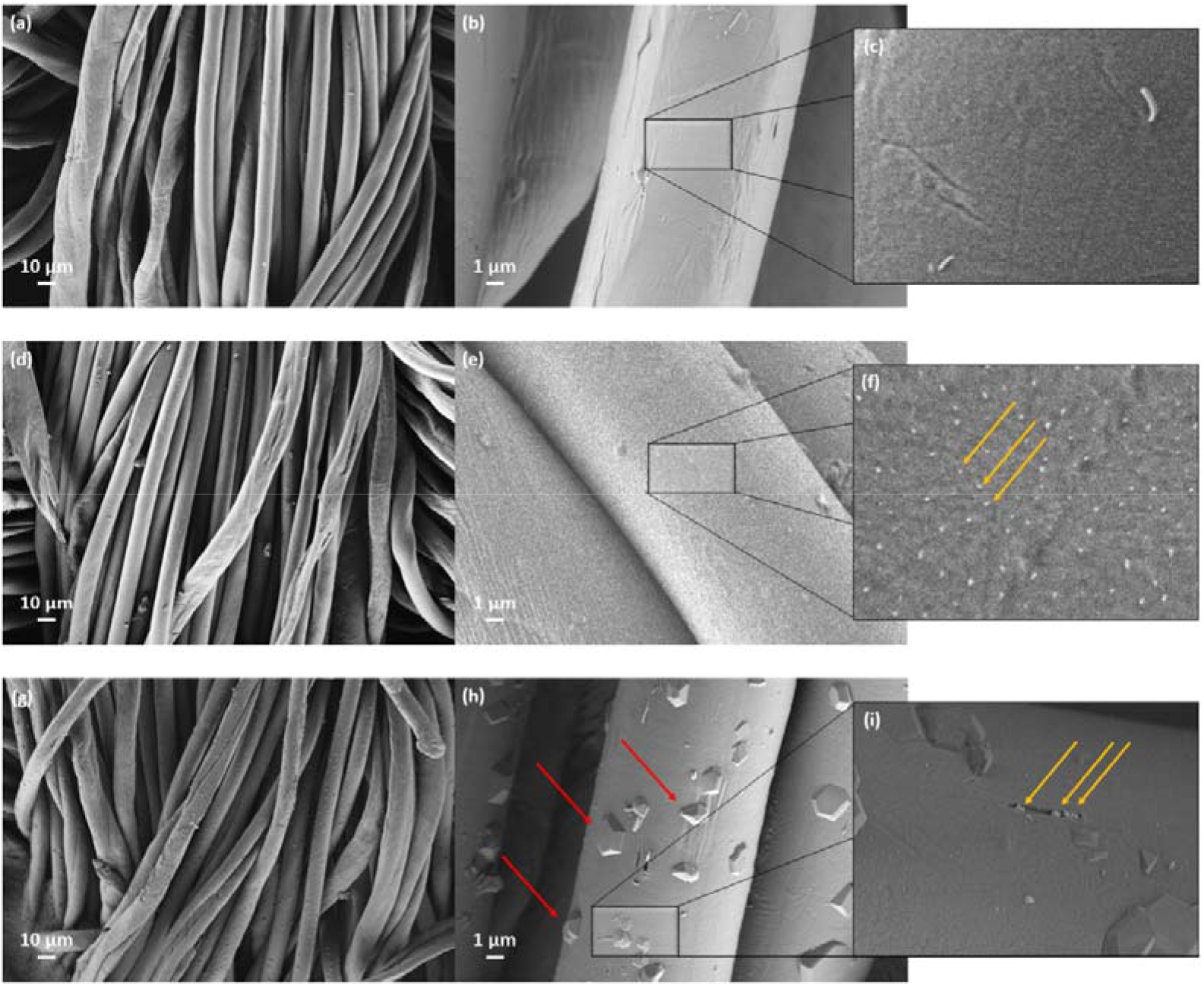
FE-SEM images of (a-c) Non-Treated Polycotton, (d-f) Polycotton AgNP-CS and (g-i) Polycotton AgNP-OP.

Since Ag-based antimicrobial additives caused distinct surface interactions in polycotton fibers, it is expected that this difference will be reflected in their physical, chemical and biological properties. In this sense, experiments were carried out to evaluate the biological properties of composites obtained through the allergenic response to humans and microbicidal activity against *E. coli*, *S. aureus, C. albicans* and SARS-CoV-2.

The AATCC 147 test results against *S. aureus* (gram positive) and *E. coli* (gram negative) bacteria for the non-treated and the Ag-based antimicrobial treated polycotton samples are shown in Figure 4 and Table 4. For the control polycotton, growth of *E. coli* and *S. aureus* was observed under the specimen while no growth appeared for the treated fabric. The zone of inhibition for the control sample was 0 mm, in comparison to 2-3 mm for the treated fabric. It can be seen from these results that the Ag-based antimicrobials treated fabrics displayed a high level of antibacterial performance.

**Table 4.**
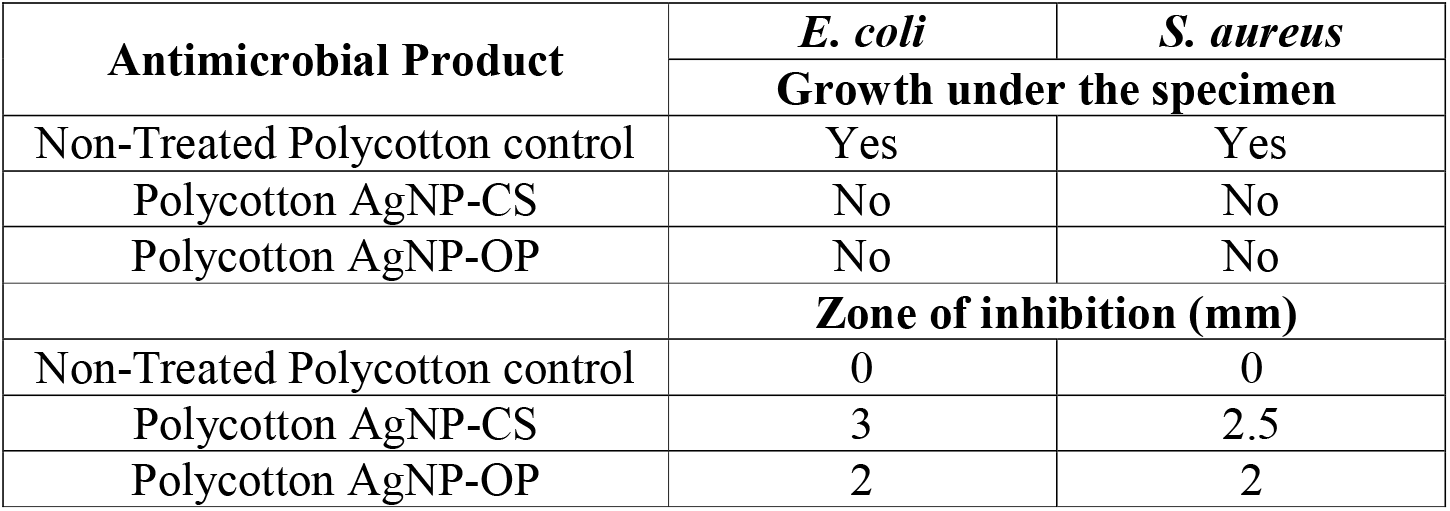
AATCC 147 tests results against *S. aureus* and *E. coli* for non-treated (control) and Ag-based antimicrobials treated fabric samples.

**Figure 4.**
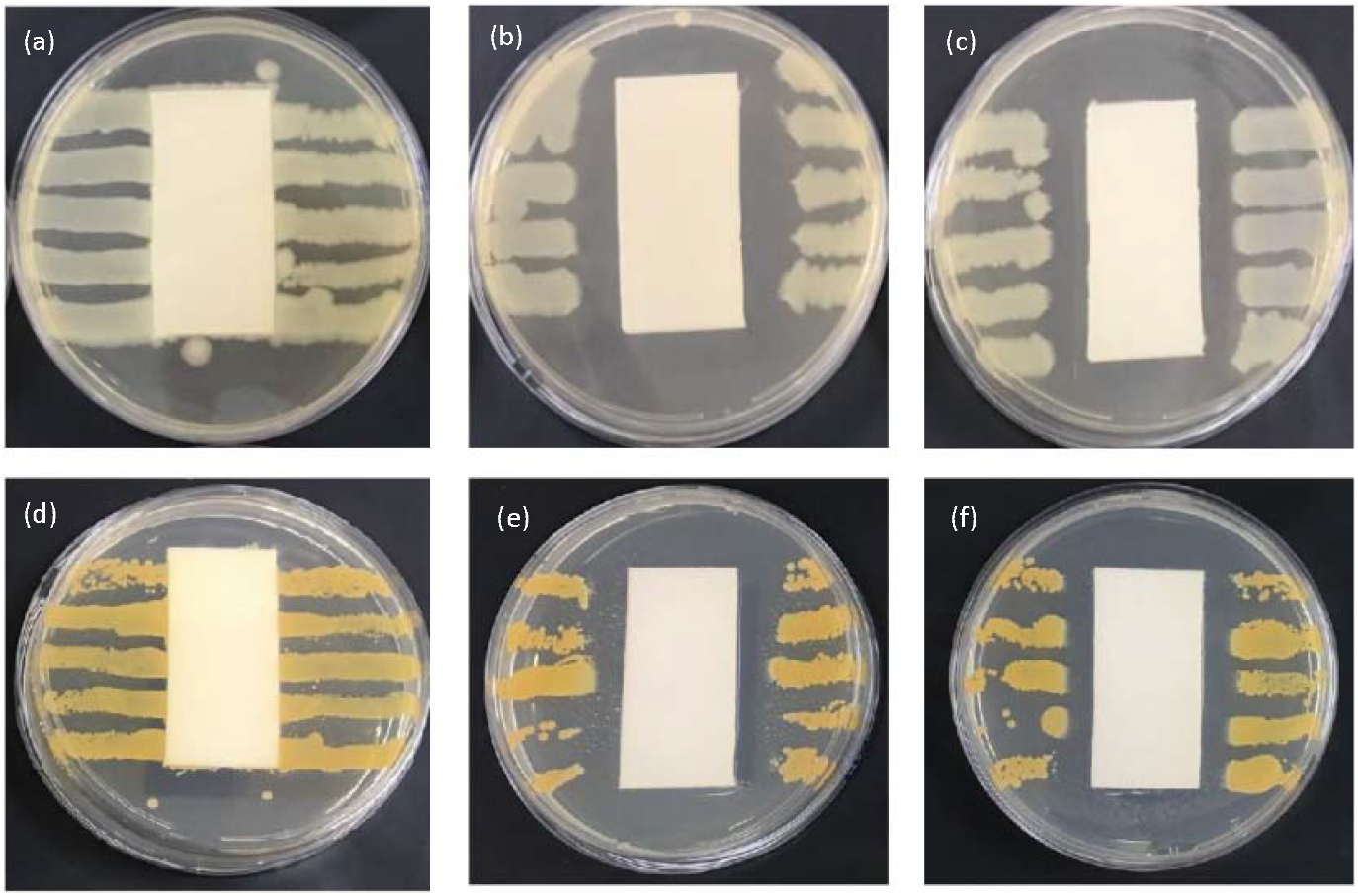
AATCC 147 test result against *E. coli* for the Non-Treated Polycotton sample as a reference (a) and for the Ag-based antimicrobial treated polycotton (b – Polycotton AgNP-CS and c – Polycotton AgNP-OP) and AATCC 147 test result against *S. aureus* for the Non-Treated Polycotton sample as a reference (d) and for the Ag-based antimicrobial treated fabric (e –Polycotton AgNP-CS and f – Polycotton AgNP-OP) exhibiting, respectively, no peripheral inhibition and a measurable zone of inhibition.

The antibacterial mechanism for gram-positive and gram-negative bacteria is associated with Ag NPs and their penetration into the cell membrane of these microorganisms. Ag NPs are able to penetrate cell membranes and release Ag^+^ ions, which have a high affinity to react with phosphorus and sulfur compounds, either from the membrane or from inside of the cell.[61] In addition, Ag NPs can generate reactive oxygen species (ROS), which cause an accumulation of intracellular ROS, leading to bacterial death from oxidative stress[62].

The halo inhibition test (AATCC 147) is only a qualitative test that shows bacterial inhibition, requiring a qualitative test to determine the percentages of inhibition (AATCC 100). The quantitative antimicrobial activities of finished textiles treated with the two different Ag-based antimicrobials according to the AATCC 100 standard are shown in Table 5. These tests were also performed with *C. albicans*, in order to assess the fungicidal potential of the Ag-based fabrics. In agreement with the qualitative test, the quantitative test showed that all the Ag-based treated polycotton samples had efficient antimicrobial activities, against bacteria and fungi, displaying a 99.99% reduction in all tested samples.

**Table 5.** Quantitative antibacterial results according to the AATCC 100 standard.

| Antimicrobial Product | Microbial Reduction<sup>a</sup> (%) |  |  |  |  |  |  |  |  |
| --- | --- | --- | --- | --- | --- | --- | --- | --- | --- |
|  | *S. aureus* ATCC 6538 |  |  | *E. coli* ATCC 8739 |  |  | *C. albicans* ATCC 10231 |  |  |
|  | Zero-time bacteria count | Bacteria count after 24 hours | % Reduction | Zero-time bacteria count | Bacteria count after 24 hours | % Reduction | Zero-time Fungi count | Fungi count after 24 hours | % Reduction |
| Non-Treated Polycotton control | $2.1 \times 10^5$ | $2.2 \times 10^5$ | - | $2.3 \times 10^5$ | $2.2 \times 10^5$ | - | $2.0 \times 10^5$ | $2.2 \times 10^5$ | - |
| Polycotton AgNP-CS | $2.1 \times 10^5$ | $1.6 \times 10$ | 99.99% | $2.3 \times 10^5$ | $1.3 \times 10$ | 99.99% | $2.0 \times 10^5$ | $1.3 \times 10$ | 99.99% |
| Polycotton AgNP-OP | $2.1 \times 10^5$ | $1.1 \times 10$ | 99.99% | $2.3 \times 10^5$ | $1.1 \times 10$ | 99.99% | $2.0 \times 10^5$ | $1.4 \times 10$ | 99.99% |

<sup>a</sup>Percent bacterial reduction as measured against a non-treated control.

The antiviral activity test was designed to determine the inactivation of viral particles upon short exposure to the products, which in this case were the Ag-based treated polycotton samples incubated in liquid media. After a short period of incubation, the media were transferred to a cell culture, where viable virions would be able to enter cells and replicate within. The supernatant of cell cultures was recovered after 48 h and the viral load was determined by RT-qPCR, resulting in the determination of number of viral RNA copies per mL.

Table 6 shows the number of copies of the control media without any fabric sample, non-treated polycotton, and the two Ag-based treated polycotton samples at the two different tested time periods. With the result of the number of copies of each sample, the viral inactivation effect of each cloth was calculated, using the media without any fabric sample as control.

**Table 6.**
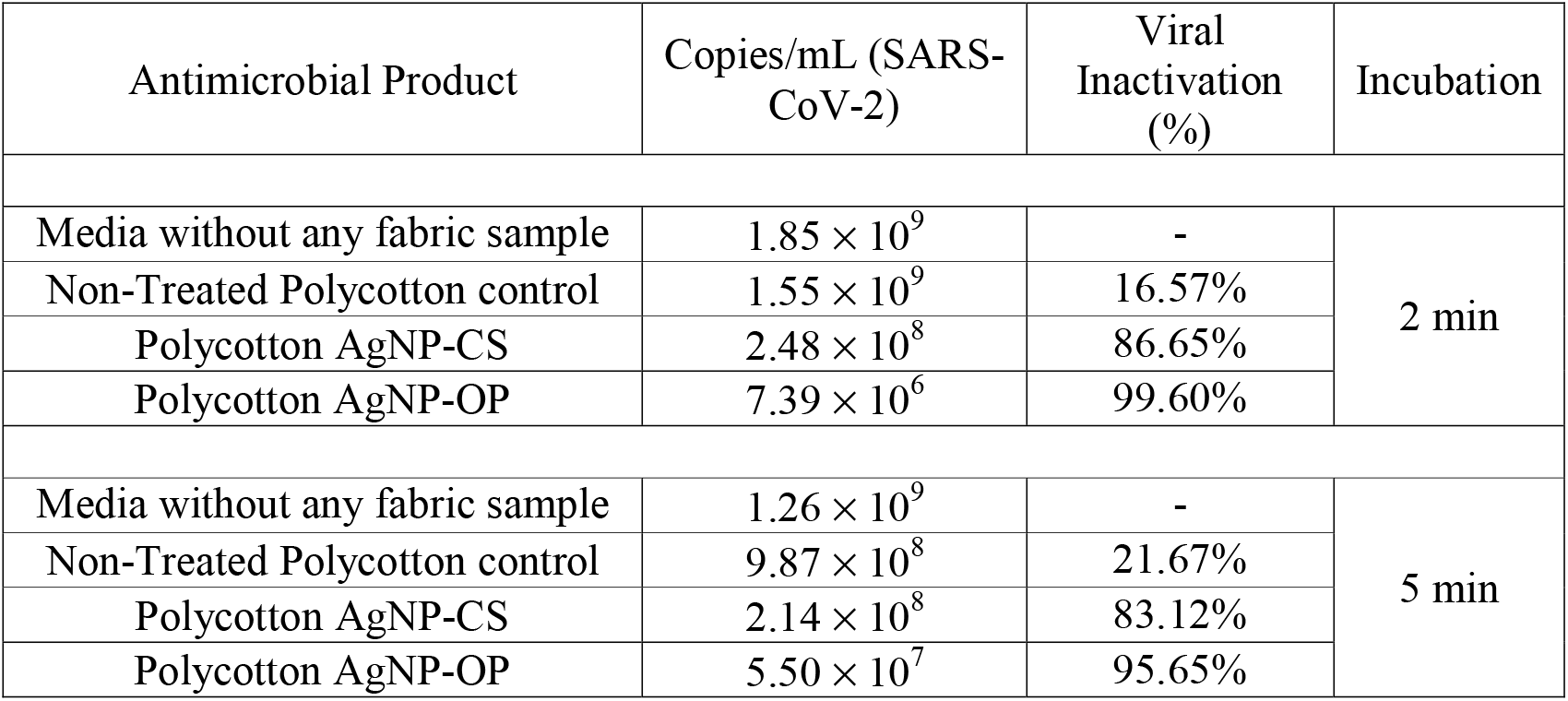
Copies per mL of SARS-CoV-2 at different times in the first experiment

| Antimicrobial Product | Copies/mL (SARS-CoV-2) | Viral Inactivation (%) | Incubation |
| --- | --- | --- | --- |
| Media without any fabric sample | $1.85 \times 10^9$ | - | 2 min |
| Non-Treated Polycotton control | $1.55 \times 10^9$ | 16.57% | |
| Polycotton AgNP-CS | $2.48 \times 10^8$ | 86.65% | |
| Polycotton AgNP-OP | $7.39 \times 10^6$ | 99.60% | |
| Media without any fabric sample | $1.26 \times 10^9$ | - | 5 min |
| Non-Treated Polycotton control | $9.87 \times 10^8$ | 21.67% | |
| Polycotton AgNP-CS | $2.14 \times 10^8$ | 83.12% | |
| Polycotton AgNP-OP | $5.50 \times 10^7$ | 95.65% | |

Regarding the second experiment, the number of copies per milliliter in each sample was also obtained and the percentage of inhibition of the products was calculated from the control media without any fabric sample. The obtained results were summarized in Table 7.

**Table 7.**
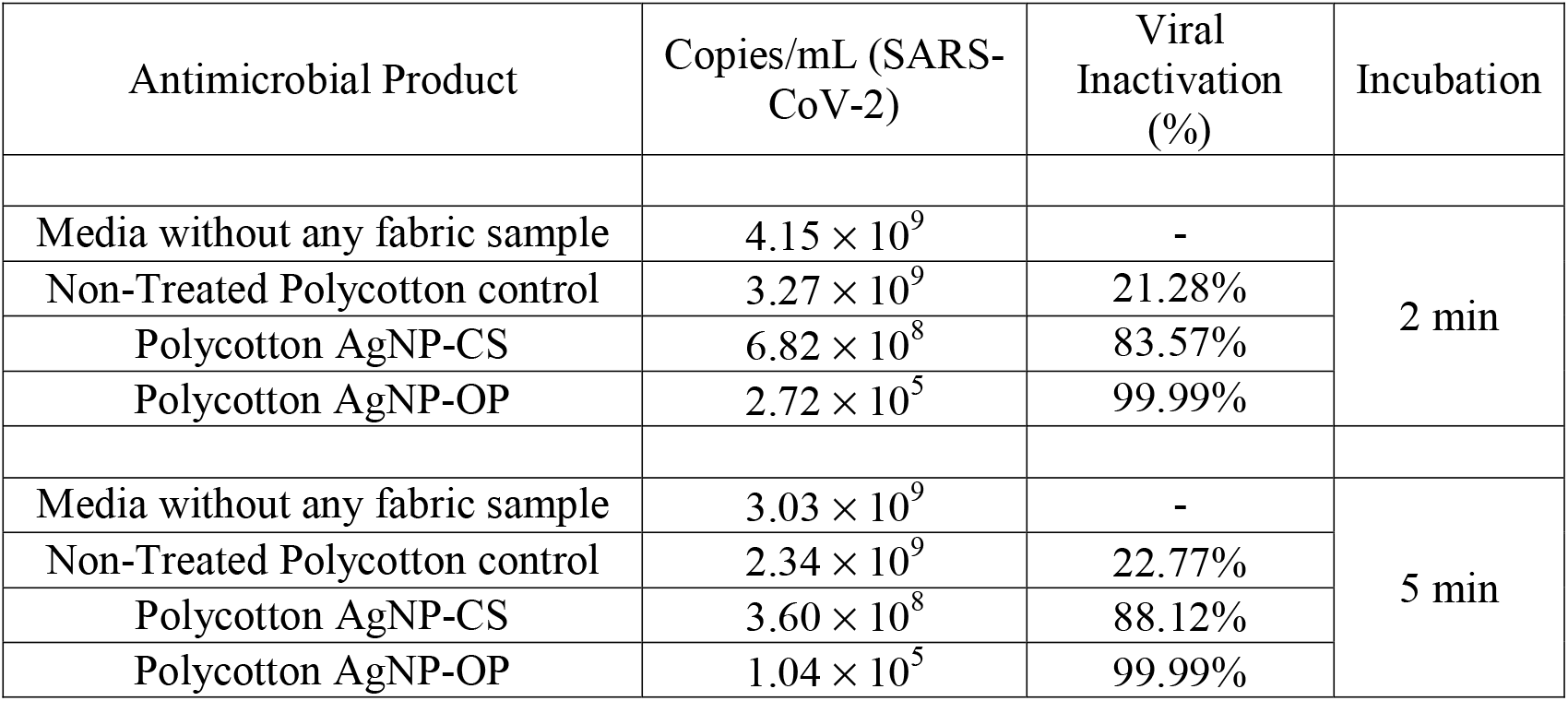
Copies per mL of SARS-CoV-2 at different times in the second experiment.

The following graphs represent the data described in Tables 5 and 6, of the control media without any fabric sample, non-treated polycotton, and the two Ag-based treated polycotton samples.

Figures 5 and 6 show the results of the first and second experiments, respectively, indicating the number of viral copies per mL and the percentage of inhibition of each compound above the bar referring to it. Inhibition was calculated for each treatment using its respective control.

**Figure 5.**
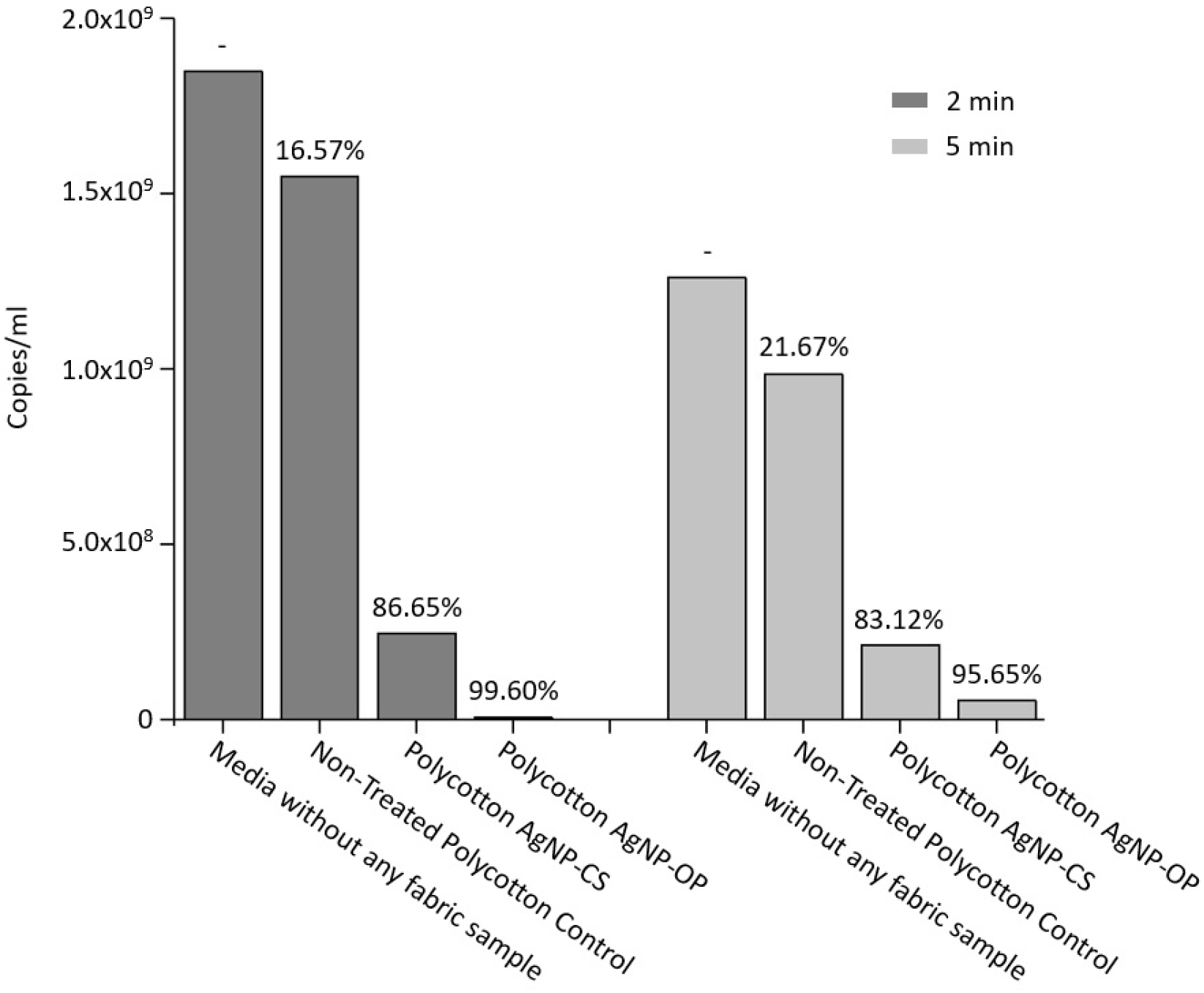
Representative graph of the data obtained in the first experiment, relating the tested products to the viral load found and the percentage of inhibition.

**Figure 6.**
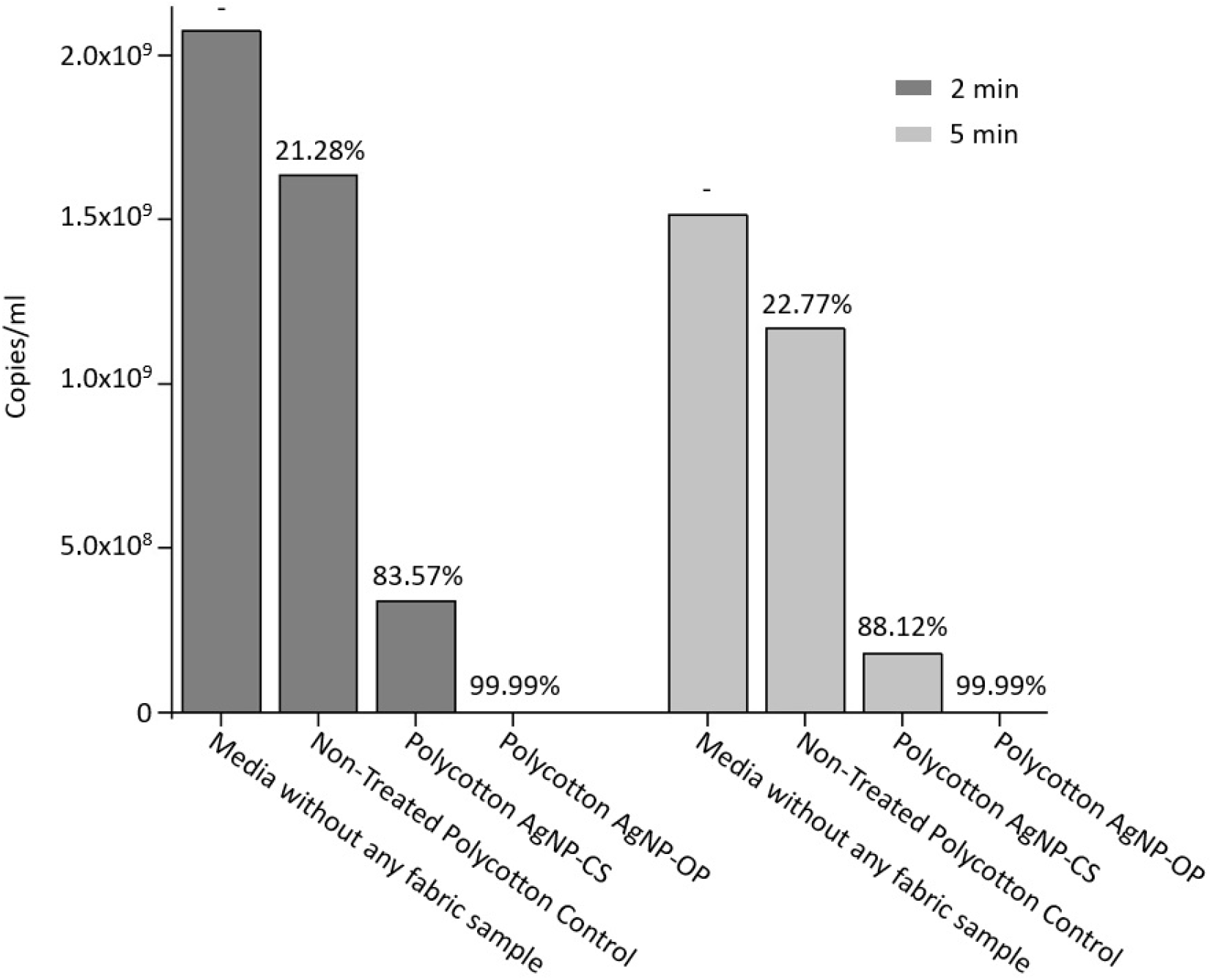
Representative graph of the data obtained in the second experiment, relating the tested products to the viral load found and the percentage of inhibition.

In both experiments, in the two time periods tested, the untreated polycotton showed a subtle activity, which was already expected by data already published by Chin and colleagues.[63] Polycotton AgNP-CS showed a high viricidal activity when incubated with the virus. At both time periods in both experiments, Polycotton AgNP-OP obtained a higher rate of viral inactivation compared to Polycotton AgNP-CS.

In short, both treated polycotton samples were effective in viral inhibition in 2 and 5 minutes in two different experiments, where there was variation in the amount of virus per cm² of fabric (4x less virus/cm² in the second experiment). Polycotton AgNP-OP showed the best activity, reaching 99.99% within two minutes of incubation with the virus in the second experiment. Polycotton AgNP-CS, despite being less effective than Polycotton AgNP-OP, showed high anti-SARS-CoV-2 activity, with more than 80% inhibition rate in all tests performed.

As an antiviral agent, Ag NPs can interfere with viral replication by two separate mechanisms of adhesion to the surface of the viral envelope. This adhesion prevents the virus from being able to connect to the infecting cell, preventing contamination and possible damage.[64,65] These antiviral mechanisms are mainly caused by stress in infected cells (due to physical contact), generation of reactive oxygen species (ROS), interactions with DNA and enzymatic damage.[66] The first mechanism is through the binding of Ag NPs with sulfur residues from the virus’s surface glycoproteins, preventing interaction with the receptor and its entry into the host cell.[67,68] The second mechanism involves the passage of Ag NPs through the cell membrane that consequently, it ends up effectively blocking the transcription factors necessary for the adequate assembly of the viral progeny.[69] Thus, in addition to the unique behavior of Ag NPs in isolation, its interface with polymers can be explored, which may open the way for new ones promising applications in several fields of action.

Both the Human Repeat Insult Patch Test and the clinical study to assess the photoirritating and photosensitizing potential were conducted according to the Cosmetic Product Safety Assessment Guide, published by the Brazilian regulatory agency ANVISA[70], by the ECOLYZER Group (São Paulo/SP, Brazil), an independent and ISO certified laboratory. For the HRIPT, Primary Dermal Irritability, Accumulated Dermal Irritability and Dermal Sensitization potential were determined. The clinical evaluation criterion was the observation of clinical signs or symptoms such as swelling (edema), redness (erythema), papules and vesicles according to the reading scale recommended by the ICDRG. No adverse reactions (erythema, edema, papules or vesicles) were detected in the product’s application areas, in the analysis of primary and accumulated irritability, sensitization, during the study period. The same clinical evaluation criterion was used to determine the Dermal Photoirritation and Dermal Photosensitization in the clinical study. As in the HRIPT, on this study no adverse reactions (erythema, edema, papules or vesicles) were detected in the product’s application areas during the study period. According to the results obtained from the sample of participants studied, we can conclude that the treated fabrics did not induce a photoirritating, photosensitizing, irritation nor sensitization process and, therefore, can be considered hypoallergenic and dermatologically tested and approved, being considered safe, according to ANVISA’s Guide for Cosmetic Product Safety.

## CONCLUSIONS

Polycotton fabrics can be functionalized to attain antibacterial, antifungal and antiviral properties using a simplistic and very common finishing treatment method in nature, the pad-dry-cure. The use of an aqueous Ag NPs solution mixed with an acrylic-based binder was demonstrated to achieve a high level of antimicrobial performance and can potentially present a high durability in relation to washing cycles as a result of the use of the binder in the impregnation solution. FTIR and Raman analyzes showed different surface effects on polycotton surface fibers due to the different chemical nature of both tested antimicrobial finishing products. Additionally, the antimicrobial finishing did not display significant differences in the fiber diameters of the samples, as shown by the FE-SEM images. Therefore, this product had excellent universality in the preparation, and it is expected that no significant change in fabric’s organoleptic properties change, requiring no special condition for its use in a major scale. Future experimental investigation would open new avenues to include the antiviral, antibacterial and antifungal treatment to a wide variety of different surfaces in addition to polycotton fabrics such as synthetic and natural fabrics, including cotton, polyesters and polyamides. Beyond the scope of this work, this simple and facile antimicrobial finishing treatment could be potentially scaled for industrial applications after addressing challenges such as choosing appropriate processing conditions and developing feasible waste disposition protocols. The main differential capability of these Ag-based fabrics is the prevention of cross infection caused by pathogens, such as opportunistic bacteria and fungi, responsible for the worsening of COVID-19 and other types of viruses. The fabrication of these fabrics composed of these materials may provide new insights into the development of protection garments and it is expected that these new textile materials may play an outstanding role as a new and important weapon against the current COVID-19 pandemic.

## Acknowledgements

The authors acknowledge the financial support of the Brazilian research financing institutions: Fundação de Amparo à Pesquisa do Estado de São Paulo (FAPESP CEPID-finance code 2013/07296◻2, Process 2017/24769-2, Process 2016/20045-7 and PIPE-finance codes 2004/08778-1 and 2017/15924-4), Coordenação de Aperfeiçoamento de Pessoal de Nível Superior - Brasil (CAPES) - Finance Code 001, Conselho Nacional de Desenvolvimento Científico e Tecnológico (CNPq) and Financiadora de Estudos e Projetos (FINEP). J. A. acknowledge Universitat Jaume I for projects UJI-B2016-25 and UJI-B2019-30, and Ministerio de Ciencia, Innovación y Universidades (Spain) project PGC2018-094417-B-I00 for supporting this research financially. We also acknowledge the Servei Informática, Universitat Jaume I for a generous allotment of computer time.

## References

[1] Carrasco-Hernandez R., Jácome R., López V. Y. and Ponce de León S. 2017 Are RNA Viruses Candidate Agents for the Next Global Pandemic? A Review. ILAR J. 58 343–58.

[2] Zhu N., Zhang D., Wang W., Li X., Yang B., Song J., Zhao X., Huang B., Shi W., Lu R., Niu P., Zhan F., Ma X., Wang D., Xu W., Wu G., Gao G. F. and Tan W. 2020 A Novel Coronavirus from Patients with Pneumonia in China, 2019. N. Engl. J. Med. 382 727–33.

[3] de Wit E., van Doremalen N., Falzarano D. and Munster V. J. 2016 SARS and MERS: recent insights into emerging coronaviruses Nat. Rev. Microbiol. 14 523–534.

[4] Sender R., Fuchs S. and Milo R. 2016 Revised Estimates for the Number of Human and Bacteria Cells in the Body PLOS Biol. 14 e1002533.

[5] Organización Mundial de la Salud 2020 Advice on the use of masks in the context of COVID-19: interim guidance-2 Guía Interna la OMS 1–5.

[6] Rai M., Yadav A. and Gade A. 2009 Silver nanoparticles as a new generation of antimicrobials Biotechnol. Adv. 27 76–83.

[7] Barillo D. J. and Marx D. E. 2014 Silver in medicine: A brief history BC 335 to present Burns 40 S3–S8.

[8] Alexander, Wesley J. 2009 History of the medical use of silver Surg. Infect. (Larchmt). 10 289–94.

[9] Vardanyan Z., Gevorkyan V., Ananyan M., Vardapetyan H. and Trchounian A. 2015 Effects of various heavy metal nanoparticles on Enterococcus hirae and Escherichia coli growth and proton-coupled membrane transport J. Nanobiotechnology 13 69.

[10] Rudramurthy G. R., Swamy M. K., Sinniah U. R. and Ghasemzadeh A. 2016 Nanoparticles: Alternatives Against Drug-Resistant Pathogenic Microbes. Molecules 21 836.

[11] Zhang P., Jiang X., Yuan P., Yan H. and Yang D. 2018 Silver nanopaste: Synthesis, reinforcements and application Int. J. Heat Mass Transf. 127 1048–69.

[12] Khatoon N., Mazumder J. A. and Sardar M. 2017 Biotechnological Applications of Green Synthesized Silver Nanoparticles J. Nanosci. Curr. Res. 02 1–2.

[13] Iravani S., Korbekandi H., Mirmohammadi S. V. and Zolfaghari B. 2014 Synthesis of silver nanoparticles: chemical, physical and biological methods Res. Pharm. Sci. 9 385–406.

[14] Abbasi E., Milani M., Fekri Aval S., Kouhi M., Akbarzadeh A., Tayefi Nasrabadi H., Nikasa P., Joo S. W., Hanifehpour Y., Nejati-Koshki K. and Samiei M. 2016 Silver nanoparticles: Synthesis methods, bio-applications and properties Crit. Rev. Microbiol. 42 173–80.

[15] Fahmy H. M., Salah Eldin R. E., Abu Serea E. S., Gomaa N. M., AboElmagd G. M., Salem S. A., Elsayed Z. A., Edrees A., Shams-Eldin E. and Shalan A. E. 2020 Advances in nanotechnology and antibacterial properties of biodegradable food packaging materials RSC Adv. 10 20467–84.

[16] Yamada M., Foote M. and Prow T. W. 2015 Therapeutic gold, silver, and platinum nanoparticles WIREs Nanomedicine and Nanobiotechnology 7 428–45.

[17] Yaqoob A. A., Umar K. and Ibrahim M. N. M. 2020 Silver nanoparticles: various methods of synthesis, size affecting factors and their potential applications–a review Appl. Nanosci. 10 1369–78.

[18] Lara H. H., Ixtepan-Turrent L., Jose Yacaman M. and Lopez-Ribot J. 2020 Inhibition of Candida auris Biofilm Formation on Medical and Environmental Surfaces by Silver Nanoparticles ACS Appl. Mater. Interfaces 12 21183–91.

[19] Gunell M., Haapanen J., Brobbey K. J., Saarinen J. J., Toivakka M., Mäkelä J. M., Huovinen P. and Eerola E. 2017 Antimicrobial characterization of silver nanoparticle-coated surfaces by “touch test” method Nanotechnol. Sci. Appl. 10 137–45.

[20] Hasan S. 2014 A Review on Nanoparticles◻: Their Synthesis and Types Res. J. Recent Sci. Res. J. Recent. Sci. Uttar Pradesh (Lucknow Campus) 4 1–3.

[21] 2006 AATCC 147-2004: Antimicrobial Activity Assessment of Textile Materials: Parallel Streak Method from American Association of Textile Chemists and Colorists

[22] 2006 AATCC 100-2004: AATCC 100:2004. Antibacterial Finishes on Textile Materials: Assessment of Developed from American Association of Textile Chemists and Colorists

[23] 20019 ISO 18184:2019. Textiles — Determination of antiviral activity of textile products.

[24] Corman V. M., Landt O., Kaiser M, Molenkamp R., Meijer A., Chu D. K., Bleicker T., Brünink S., Schneider J., Schmidt M. L., Mulders D. G., Haagmans B. L., van der Veer B., van den Brink S., Wijsman L., Goderski G., Romette J-L., Ellis J., Zambon M., Peiris M., Goossens H., Reusken C., Koopmans M. P. and Drosten C. 2020 Detection of 2019 novel coronavirus (2019-nCoV) by real-time RT-PCR. Euro Surveill. 25 20200409c.

[25] Drosten C., Günther S., Preiser W., van der Werf S., Brodt H-R., Becker S., Rabenau H., Panning M., Kolesnikova L., Fouchier R. A. M., Berger A., Burguière A-M., Cinatl J., Eickmann M., Escriou N., Grywna K., Kramme S., Manuguerra J-C., Müller S, Rickerts V., Stürmer M., Vieth S., Klenk H-D., Osterhaus A. D. M. E., Schmitz H. and Doerr H. W. 2003 Identification of a novel coronavirus in patients with severe acute respiratory syndrome. N. Engl. J. Med. 348 1967–1976.

[26] Wilkinson D. S., Fregert S., Magnusson B., Bandmann H. J., Calnan C. D., Cronin E., Hjorth N., Maibach H. J., Malalten K. E., Meneghini C. L. and Pirilä V. 1970 Terminology of contact dermatitis. Acta Derm. Venereol. 50 287–92.

[27] Fitzpatrick T. B. 1988 The Validity and Practicality of Sun-Reactive Skin Types I Through VI Arch. Dermatol. 124 869–71.

[28] Tessier D. 2013 Surface modification of biotextiles for medical applications Woodhead Publishing Series in Textiles ed M. W. King, B. S. Gupta and R. B. T-B. as M. I. Guidoin (Woodhead Publishing) pp 137–156.

[29] Moiz A., Padhye R. and Wang X. 2017 Coating of TPU-PDMS-TMS on polycotton fabrics for versatile protection Polymers (Basel). 9 660.

[30] Ran J., He M., Li W., Cheng D. and Wang X. 2018 Growing ZnO nanoparticles on polydopamine-templated cotton fabrics for durable antimicrobial activity and UV protection Polymers (Basel). 10 495.

[31] Dhineshbabu N. R. and Bose S. 2019 UV resistant and fire retardant properties in fabrics coated with polymer based nanocomposites derived from sustainable and natural resources for protective clothing application Compos. Part B Eng. 172 555–563.

[32] Liu G., Xiang J., Xia Q., Li K., Yan H. and Yu L. 2020 Fabrication of Durably Antibacterial Cotton Fabrics by Robust and Uniform Immobilization of Silver Nanoparticles via Mussel-Inspired Polydopamine/Polyethyleneimine Coating Ind. Eng. Chem. Res. 59 9666–9678.

[33] Noorian S. A., Hemmatinejad N. and Navarro J. A. R. 2020 Ligand modified cellulose fabrics as support of zinc oxide nanoparticles for UV protection and antimicrobial activities Int. J. Biol. Macromol. 154 1215–26.

[34] Joshi M and Butola B. S. 2013 Application technologies for coating, lamination and finishing of technical textiles Woodhead Publishing Series in Textiles ed M. L. B. T-A. in the D. and F. of T. T. Gulrajani (Woodhead Publishing) pp 355–411.

[35] Roy Choudhury A. K. 2017 Introduction to finishing Woodhead Publishing Series in Textiles ed A. K. B. T-P. of T. F. Roy Choudhury (Woodhead Publishing) pp 1–19.

[36] Rygula A., Jekiel K., Szostak-Kot J., Wrobel T. P. and Baranska M. 2011 Application of FT-Raman spectroscopy for in situdetection of microorganisms on the surface of textiles J. Environ. Monit. 13 2983–2987.

[37] Cabrales L., Abidi N. and Manciu F. 2014 Characterization of developing cotton fibers by confocal Raman microscopy Fibers 2 285–294.

[38] Liu Y. 1998 Vibrational spectroscopic investigation of Australian cotton cellulose fibres Part 1. A Fourier transform Raman study Analyst 123 633–636.

[39] Was-Gubala J. and Machnowski W. 2014 Application of Raman Spectroscopy for Differentiation Among Cotton and Viscose Fibers Dyed with Several Dye Classes Spectrosc. Lett. 47 527–535.

[40] Adebajo M. O., Frost R. L., Kloprogge J. T. and Kokot S. 2006 Raman spectroscopic investigation of acetylation of raw cotton Spectrochim. Acta Part A Mol. Biomol. Spectrosc. 64 448–453.

[41] Rebollar E., Pérez S., Hernández M., Domingo C., Martín M., Ezquerra T. A., García-Ruiz J. P. and Castillejo M. 2014 Physicochemical modifications accompanying UV laser induced surface structures on poly(ethylene terephthalate) and their effect on adhesion of mesenchymal cells Phys. Chem. Chem. Phys. 16 17551–9.

[42] Bauer A. J. R. 2018 Raman Spectroscopy for Fiber Analysis 023 3–6.

[43] Lin C-C., Krommenhoek P. J., Watson S. S. and Gu X. 2014 Chemical depth profiling of photovoltaic backsheets after accelerated laboratory weathering Reliab. Photovolt. Cells, Modul. Components, Syst. VII 9179 91790R.

[44] Xu L-L., Guo M-X., Liu S. and Bian S-W. 2015 Graphene/cotton composite fabrics as flexible electrode materials for electrochemical capacitors RSC Adv. 5 25244–9.

[45] Fang L., Zhang X., Ma J., Sun D., Zhang B. and Luan J. 2015 Eco-friendly cationic modification of cotton fabrics for improving utilization of reactive dyes RSC Adv. 5 45654–61.

[46] Chung C., Lee M. and Choe E. K. 2004 Characterization of cotton fabric scouring by FT-IR ATR spectroscopy Carbohydr. Polym. 58 417–20.

[47] Borazan A. A. and Gokdai D. 2018 Pine Cone and Boron Compounds Effect as Reinforcement on Mechanical and Flammability Properties of Polyester Composites Open Chem. 16 427–36.

[48] Portella E. H., Romanzini D., Angrizani C. C., Amico S. C. and Zattera A. J. 2016 Influence of Stacking Sequence on the Mechanical and Dynamic Mechanical Properties of Cotton/Glass Fiber Reinforced Polyester Composites Mater. Res. 19 542–7.

[49] Al-Balakocy N., El-Badry K. and Hassan T. 2019 Multi-Finishing of Polyester and Polyester Cotton Blend Fabrics Activated by Enzymatic Treatment and Loaded with Zinc Oxide Nanoparticles.InterchOpen 1 1–14.

[50] Xu Q., Xie L., Diao H., Li F., Zhang Y., Fu F. and Liu X. 2017 Antibacterial cotton fabric with enhanced durability prepared using silver nanoparticles and carboxymethyl chitosan Carbohydr. Polym. 177 187–193.

[51] Yan L., Hao Y., Feng X., Yang Y., Liu X., Chen Y. and Xu B. 2015 Synthesis and optical properties of composite films from P3HT and sandwich-like Ag–C– Ag nanoparticles RSC Adv. 5 79860–7.

[52] Liu H., Lv M., Deng B., Li J., Yu M., Huang Q. and Fan C. 2014 Laundering durable antibacterial cotton fabrics grafted with pomegranate-shaped polymer wrapped in silver nanoparticle aggregations Sci. Rep. 4 5920.

[53] Nam S., Condon B. D., Delhom C. D. and Fontenot K. R. 2016 Silver-cotton nanocomposites: Nano-design of microfibrillar structure causes morphological changes and increased tenacity Sci. Rep. 6 37320.

[54] Xu Q., Li R., Shen L., Xu W., Wang J., Jiang Q., Zhang L., Fu F., Fu Y. and Liu 2019 Enhancing the surface affinity with silver nano-particles for antibacterial cotton fabric by coating carboxymethyl chitosan and l-cysteine Appl. Surf. Sci. 497 143673.

[55] Zhang.F, Wu X., Chen Y. and Lin H. 2009 Application of silver nanoparticles to cotton fabric as an antibacterial textile finish Fibers Polym. 10 496–501.

[56] Chen C-Y. and Chiang C-L. 2008 Preparation of cotton fibers with antibacterial silver nanoparticles Mater. Lett. 62 3607–9.

[57] Maráková N., Humpolíček P., Kašpárková V., Capáková Z., Martinková L., Bober P., Trchová M. and Stejskal J. 2017 Antimicrobial activity and cytotoxicity of cotton fabric coated with conducting polymers, polyaniline or polypyrrole, and with deposited silver nanoparticles Appl. Surf. Sci. 396 169–176.

[58] Ghosh S., Yadav S. and Reynolds N. 2010 Antibacterial properties of cotton fabric treated with silver nanoparticles J. Text. Inst. 101 917–924.

[59] Montazer M., Alimohammadi F., Shamei A. and Rahimi M. K. 2012 Durable antibacterial and cross-linking cotton with colloidal silver nanoparticles and butane tetracarboxylic acid without yellowing Colloids Surfaces B Biointerfaces 89 196–202.

[60] Bacciarelli-Ulacha A., Rybicki E., Matyjas-Zgondek E., Pawlaczyk A. and Szynkowska M. I. 2014 A New Method of Finishing of Cotton Fabric by in Situ Synthesis of Silver Nanoparticles Ind. Eng. Chem. Res. 53 4147–55.

[61] Gao M., Sun L., Wang Z. and Zhao Y. 2013 Controlled synthesis of Ag nanoparticles with different morphologies and their antibacterial properties Mater. Sci. Eng. C 33 397–404.

[62] Assis M., Robeldo T., Foggi C. C., Kubo A. M., Mínguez-Vega G., Condoncillo E., Beltran-Mir H., Torres-Mendieta R., Andrés J., Oliva M., Vergani C. E., Barbugli P. A., Camargo E. R., Borra R. C. and Longo E. 2019 Ag Nanoparticles/α-Ag_2_WO_4_ Composite Formed by Electron Beam and Femtosecond Irradiation as Potent Antifungal and Antitumor Agents Sci. Rep. 9 9927.

[63] Chin A. W. H., Chu J. T. S., Perera M. R. A., Hui K. P. Y., Yen H-L., Chan M. C. W., Peiris M. and Poon L. L. M. 2020 Stability of SARS-CoV-2 in different environmental conditions The Lancet Microbe 5247 2004973.

[64] Alkhouri N. and Zein N. N. 2012 Protease inhibitors: Silver bullets for chronic hepatitis C infection? Cleve. Clin. J. Med. 79 213–22.

[65] Khylko O. L. 2016 Possible mechanism of inhibition of virus infectivity with nanoparticles Semicond. Phys. Quantum Electron. Optoelectron. 19 220–224.

[66] Akbarzadeh A., Kafshdooz L., Razban Z., Dastranj Tbrizi A., Rasoulpour S., Khalilov R., Kavetskyy T., Saghfi S., Nasibova A. N., Kaamyabi S. and Kafshdooz T. 2018 An overview application of silver nanoparticles in inhibition of herpes simplex virus Artif. Cells, Nanomedicine, Biotechnol. 46 263–726.

[67] Speshock J. .L, Murdock R. C., Braydich-Stolle L. K., Schrand A. M. and Hussain S. M. 2010 Interaction of silver nanoparticles with Tacaribe virus J. Nanobiotechnology 8 19.

[68] Lara H. H., Ayala-Nuñez N. V., Ixtepan-Turrent L. and Rodriguez-Padilla C. 2010 Mode of antiviral action of silver nanoparticles against HIV-1 J. Nanobiotechnology 8 1.

[69] Greulich C., Diendorf J., Simon T., Eggeler G., Epple M. and Köller M. 2011 Uptake and intracellular distribution of silver nanoparticles in human mesenchymal stem cells Acta Biomater. 7 347–54.

[70] Agência Nacional de Vigilânia Sanitária 2012 Guia para Avaliação de Segurança de Produtos Cosméticos Guia para Avaliação de Segurança de Produtos Cosméticos Anvisa 2 1–74.

